# Semaphorin 3G exacerbates joint inflammation through the accumulation and proliferation of macrophages in the synovium

**DOI:** 10.1101/2022.02.12.480222

**Authors:** Jumpei Shoda, Shigeru Tanaka, Keishi Etori, Koto Hattori, Tadamichi Kasuya, Kei Ikeda, Yuko Maezawa, Akira Suto, Kotaro Suzuki, Junichi Nakamura, Yoshiro Maezawa, Minoru Takemoto, Christer Betsholtz, Koutaro Yokote, Seiji Ohtori, Hiroshi Nakajima

**Affiliations:** Department of Allergy and Clinical Immunology, Graduate School of Medicine, Chiba University, Chiba, Japan; Department of Orthopedic Surgery, Graduate School of Medicine, Chiba University, Chiba, Japan; Department of Endocrinology, Hematology, and Gerontology, Graduate School of Medicine, Chiba University, Chiba, Japan; Department of Medicine, Division of Diabetes, Metabolism and Endocrinology, International University of Health and Welfare, Narita, Japan; Department of Immunology, Genetics and Pathology (IGP), Uppsala University, Uppsala, Sweden

**Keywords:** Rheumatoid arthritis, Semaphorin, Macrophage, Neuropilin-2, Methotrexate

## Abstract

**Objective:** Methotrexate (MTX) is an anchor drug for rheumatoid arthritis (RA) treatment; however, the exact mechanisms by which MTX improves RA activity are still debatable. This study aimed to understand the roles of molecules whose expression is affected by MTX in RA patients and find novel therapeutic targets.

**Methods:** CD4^+^ T cells from 28 treatment naïve RA patients before and 3 months after the initiation of MTX treatment were subjected to DNA microarray analyses. The expression of Semaphorin 3G (Sema3G), as one of the differentially-expressed genes, and its receptor, Neuropilin-2 (Nrp2), was evaluated in RA synovium and collagen-induced arthritis (CIA) synovium. CIA and collagen antibody-induced arthritis (CAIA) were induced in Sema3G-deficient (Sema3G^-/-^) mice and control mice, and the clinical score, histological score, and serum cytokines were assessed. The migration and proliferation of Sema3G-stimulated bone marrow-derived macrophages (BMMs) were analyzed in vitro. The effect of local Sema3G administration during CAIA on the clinical score and the quantity of infiltrating macrophages was evaluated.

**Results:** The expression of Sema3G in CD4^+^ T cells was downregulated by MTX treatment in RA patients. Sema3G was expressed in RA but not osteoarthritis synovium, and its receptor Nrp2 was mainly expressed on activated macrophages. Sema3G deficiency ameliorated CIA and CAIA. Sema3G stimulation enhanced the migration and proliferation of BMMs. The local administration of Sema3G deteriorated CAIA and increased infiltrating macrophages.

**Conclusions:** Upregulation of Sema3G in RA synovium is a novel mechanism to deteriorate joint inflammation through the accumulation of macrophages.

**Key messages:** Semaphorin 3G is expressed in the inflamed synovium in human and mice.

The receptor of Semaphorin 3G is mainly expressed on M1 macrophages.

Semaphorin 3G deteriorates inflammatory arthritis through macrophage proliferation and migration.

## Introduction

Rheumatoid arthritis (RA) is an autoimmune disease characterized by chronic destructive synovial inflammation. Recent advances in RA treatment have improved the outcome. However, there are substantial numbers of patients who do not achieve remission even when biologic disease-modifying anti-rheumatic drugs (biologic DMARDs, bDMARDs) or target synthetic DMARDs (tsDMARDs) accompanied with methotrexate (MTX) were used [1]. Also, these drugs increase the risk of serious infections [2, 3]. This is partly because the molecular targets of bDMARDs, tsDMARDs, and MTX are highly involved in the global pro-inflammatory pathways. In this regard, it is crucial to narrow down target pathways to reduce the adverse event while guaranteeing the efficacy of RA.

MTX is an anchor drug for RA treatment. MTX is a folate derivative that inhibits nucleotide synthesis, resulting in decreased immune cell proliferation [4]. In addition to this function, several studies revealed the other mechanisms by which MTX suppresses the immune responses [5–8]; however, it is required to understand more about the pharmacological action of MTX because of the point mentioned above. Thus, we aimed to analyze the molecular mechanisms improving RA pathology by MTX and identify novel therapeutic targets.

Here, we examined gene expression profiles of peripheral blood CD4^+^ T cells in 28 treatment-naïve RA patients before and after MTX treatment and evaluated the roles of one of the differentially-expressed genes, Semaphorin 3G (Sema3G), in murine experimental arthritis models.

## Materials and Methods

### Patients

Twenty-eight patients who fulfilled the American College of Rheumatology 1987 revised criteria for the classification of RA and who never received any DMARDs were recruited for DNA microarray analysis. The patients started MTX treatment as standard clinical care. Information on the patient characteristics was described in Table 1. For Sema3G detection, 8 RA and 8 osteoarthritides (OA) patients who received joint surgery were recruited, and the synovium was trimmed from the surgical specimen. The procedure was approved by the Ethics Committee of Chiba University (reference number 872 and 3880), and written informed consent was obtained in accordance with the Declaration of Helsinki.

**Table 1.** Patient characteristics.

| Variables | Treatment naïve RA (n = 28) |
| --- | --- |
| Age, median (IQR) years | 59.5 (47.5 - 67) |
| Female, n (%) | 20 (71.4) |
| Disease duration, median (IQR) months | 7 (3 - 24) |
| RF positive, n (%) | 25 (89.2) |
| ACPA positive, n (%) | 26 (92.9) |
| Maximum dose of MTX, median (IQR) mg/week | 10 (8 - 11) |
| Dose of prednisolone at baseline, median (IQR) mg/day | 0 (0 - 0) |
| ESR, median (IQR) mm/hour Pre-treatment | 31 (14 - 46) |
| ESR, median (IQR) mm/hour Post-treatment | 11 (9 - 27) |
| CRP, median (IQR) mg/dl Pre-treatment | 0.7 (0.3 - 1.5) |
| CRP, median (IQR) mg/dl Post-treatment | 0.1 (0.1 - 0.4) |
| DAS28-ESR, median (IQR) Pre-treatment | 4.21 (3.83 - 5.02) |
| DAS28-ESR, median (IQR) Post-treatment | 2.54 (2.02 - 4.39) |

IQR, inter-quartile range; RF, Rheumatoid factor; ACPA, anti-cyclic citrullinated protein antibody; MTX, methotrexate; ESR, erythrocyte sedimentation rate; CRP, C-reactive protein; DAS28, Disease Activity Score 28

### Mice

C57BL6/J mice were purchased from Japan Clea. Sema3G-deficient (Sema3G^-/-^) mice were described previously [9]. Mice were housed in specific pathogen-free facilities. All animal experiments were conducted in accordance with the Animal Care and Use Committee at Chiba University.

### Collagen-induced arthritis and collagen antibody-induced arthritis

For collagen-induced arthritis (CIA), 8-12-week-old mice were immunized intradermally at the base of the tail with 100 μl of chicken type II collagen (10 μg/ml) (Chondrex) emulsified with complete Freund’s adjuvant (20 μg/ml Mycobacterium butyricum) (Chondrex). Three weeks later, mice received the same immunization. After the second immunization, the clinical scores were recorded 3 times a week. For collagen antibody-induced arthritis (CAIA), 8-12-week-old mice received an intravenous injection of 100 μl ArthritoMab (MD Bioproducts). On day 4, mice were injected with 100 μg LPS intraperitoneally. The clinical score was recorded daily. The clinical symptom was graded based on the criteria as follows. 0 = no joint swelling; 1 = mild joint swelling/ erythema of ankle, wrist, or digit; 2 = moderate joint swelling; 3 = severe joint swelling; 4 = severe joint swelling with ankylosis. The clinical score was obtained by summing the scores for each limb.

### Antibodies

Anti-Sema3G polyclonal antibody was from Mybiosource. Anti-Neuropilin2 (Nrp2) polyclonal antibody was from Alomone Labs. Antibodies against CD45 (104), CD3ε (145-2C11), CD4 (GK1.5), B220 (RA3-6B2), CD11b (M1/70), F4/80 (BM8), CD86 (GL-1), and MHC-II (M5/114.15.2), and Zombie Aqua were from BioLegend.

### Recombinant Fc-tagged Sema3G production

Sema3G expression vector was described elsewhere [10]. The vector was transfected into 293T cells using Lipofectamine LTX (Thermo Fisher Scientific). Cells were cultured in DMEM supplemented with ultra-low IgG FBS (Gibco). Culture supernatant was collected and dialyzed using Amicon Ultra 100K (Merck). Dialyzed supernatant was further purified with Protein G column (Cytiva), and then the buffer was extensively exchanged to PBS. Production of Sema3G was confirmed by Coomassie blue staining and western blotting.

### Immunohistochemistry

Formalin-fixed paraffin-embedded samples were subjected to Sema3G immunostaining. After dewaxing and rehydration, the samples were boiled in 0.01M citric acid buffer (pH6.0) for 10 minutes. The samples were then washed and blocked in 3% BSA solution followed by overnight primary antibody reaction (antibody dilution rate = 1:50). After washing, the samples were subjected to secondary antibody staining and visualization with DAB using Liquid DAB Substrate/Chromogen System (DACO).

### Bone marrow-derived macrophage culture

Bone marrow cells were cultured in the presence of recombinant M-CSF (50 ng/ml) (Pepro Tech) to prepare bone marrow-derived macrophages (BMMs). For Nrp2 expression analysis, BMMs were stimulated with either 100 ng/ml LPS (Sigma Aldrich), 10 ng/ml IFNγ (Pepro Tech), or 20 ng/ml IL-4 (Pepro Tech) for 2 days. For RNA-seq analyses, LPS-stimulated BMMs were cultured in the presence or absence of 100 ng/ml recombinant Sema3G for 18 hours. For cell proliferation analyses, 5×10^4^ BMMs were seeded in a 24-well plate and cultured with or without recombinant Sema3G for 48 hours. Cells were incubated with EdU for the last 2 hours, followed by Hoechst 33342 and EdU staining (Click-iT EdU assay, Thermo Fisher Scientific).

### Transwell migration assay

Transwell migration assay was performed as previously described [11]. Briefly, upper chambers with 8 μm pore size membranes were filled with LPS-stimulated BMMs in a 1% FBS supplemented medium. The lower chambers were filled with 700 mL of 1% FBS supplemented medium with PBS, Sema3G, or MCP1 (BioLegend) as indicated. After 6-hour stimulation, cells on the underside of the membrane were stained with Hoechst.

### Flow cytometry

Joint infiltrating cells were isolated as previously described [12]. Cells were first stained with Zombie Aqua to exclude dead cells from the analyses and subsequently incubated with anti-CD16/32 antibody (BioLegend) to block Fc receptors. Samples were then stained with antibodies against CD45, CD3ε, CD4, B220, CD11b, F4/80, and Nrp2 at 4°C for 30 minutes. For the analyses of BMMs, adhered cells were detached using 5mM EDTA in PBS and then stained with anti-Nrp2 antibody. Samples were analyzed on a Canto II or a Fortessa X-20 (BD Biosciences).

### Microarray analysis

Peripheral blood samples were prepared from MTX-treated patients as previously described [13], and CD4^+^ T cells were further enriched by using a human CD4^+^ T cell isolation kit (Miltenyi) [14–16]. Total RNA was extracted using Isogen solution (Nippon Gene). DNA microarray analysis was performed using a Quick Amp labeling kit and a Whole Human Genome DNA Microarray 4×44K according to the manufacturer’s protocol (Agilent). Signal intensity was normalized by adjusting the data to a 75th percentile value. A Linear Models for Microarray Data (Limma) package of R project was used to identify candidate probes [17].

### RNA-seq analysis

Total RNA was purified with a Trizol reagent. RNA-seq libraries were prepared using a QuantSeq 3’ mRNA-Seq Library Prep Kit FWD for Illumina and UMI Second Strand Synthesis Module for QuantSeq FWD (Lexogen). Sequencing was performed on an Illumina NexSeq500 (Illumina) in a 75-base single-end mode. Obtained reads were mapped on mm10 genome, and UMI counts were measured using Strand NGS (Agilent). Expression data were normalized, and differentially expressed genes were identified with the EdgeR.

### ELISA

Serum levels of IL-6 and TNFα were measured by a mouse ELISA MAX Standard IL-6 kit (BioLegend) and a mouse ELISA MAX Standard TNFα kit (BioLegend).

### Statistical analyses

An unpaired, paired t-test, or one-way ANOVA test was used for data analysis on Prism9 (GraphPad). The clinical score was assessed by a two-way ANOVA test. P values < 0.05 were considered significant.

### Data availability

Microarray and RNA-seq data have been deposited in Gene Expression Omnibus and are accessible through GSE176440 and GSE176438. The data will be publicly available upon publication.

## Results

### Semaphorin 3G expression is increased in the inflamed joint

We first performed DNA microarray analyses of peripheral blood CD4^+^ T cells before and after MTX treatment to find novel mechanisms by which MTX improves RA disease activity. We identified several differentially-expressed genes and focused on Sema3G (Figure 1A). Sema3G belongs to the class 3 semaphorin family, and it has been revealed that Sema3G has roles in neural and vascular development [18, 19]. Although several reports have suggested important roles of semaphorins in autoimmune diseases [20], the role of Sema3G in this context is yet to be elucidated. Therefore, we analyzed Sema3G expression in human synovium obtained from RA and OA patients. As described previously, OA synovium was monolayer, and the synoviocytes and fibroblast-like spindle-shaped cells were slightly positive for Sema3G (Figure 1B). In contrast, multi-layered synoviocytes were observed in RA synovium and expressed substantial levels of Sema3G (Figure 1B). In addition, synovium-infiltrating leukocytes expressed Sema3G (Figure 1B). We compared the Sema3G-positive area in the synovial tissue and found that the Sema3G-positive area was significantly larger in RA synovium than in OA synovium (Figure 1C).

**Figure 1.**
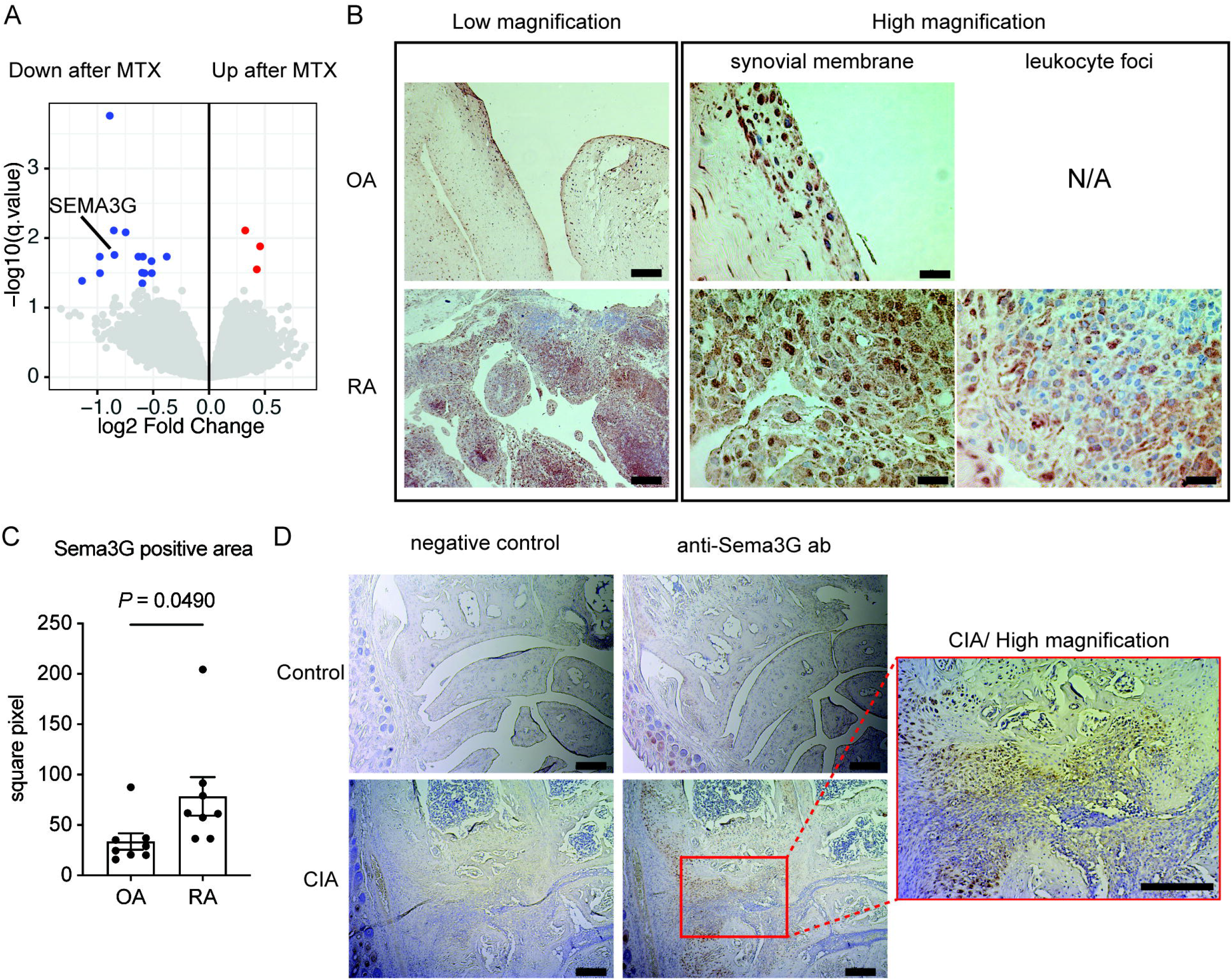
The enhanced expression of Sema3G in inflamed synovium in humans and mice. **A.** Genes differentially expressed before and after MTX treatment. Genes differentially expressed are highlighted in red (up after MTX) or blue (down after MTX). **B.** Sema3G expression in the synovium of OA or RA patients. The synovium specimens were stained with anti-Sema3G antibody and visualized with DAB. Bars indicate 200 μm (Low magnification) or 20 μm (high magnification). **C.** The cumulative data of Sema3G-positive area. Data are expressed means ± SEM. The statistical analyses were performed using an unpaired t-test. **D.** Sema3G expression in the synovium of CIA. The hind paw of control and CIA-induced mice were stained with anti-Sema3G antibody. The representative data are shown. Similar results were obtained in 3 independent experiments. Bars indicate 200 μm.

We next assessed Sema3G expression in CIA. Non-arthritic control mice have minimum synovium in wrist joints with no Sema3G signals (Figure 1D). Meanwhile, CIA-induced mice showed thick synovium with abundant Sema3G signals (Figure 1D). These results implicate that Sema3G has some roles in the pathogenesis of joint inflammation.

### Activated macrophages express Neuropilin-2, a receptor for Sema3G, in the inflamed joint

Previous reports showed that Nrp2 and plexin complex [21] functions as the receptor for full-length Sema3G. To clarify the cell types that respond to Sema3G in the inflamed joint, joint-infiltrating immune cells were collected from CIA-induced mice and analyzed for the expression of Nrp2. Among several immune cells, macrophages showed the highest percentage of the Nrp2-positive population (Figure 2A). Of note, Nrp2-positive macrophages expressed higher levels of CD86 and MHC class II, which are representative activation markers (Figure 2B), suggesting that the signals through Nrp2 might be involved in the activation of macrophages. Next, BMMs were cultured under several conditions to understand what type of macrophages preferentially expresses Nrp2. Without any stimulations, only 20% of BMMs were positive for Nrp2 (Figure 2C). Importantly, IFNγ or LPS stimulation efficiently induced Nrp2 expression, but IL-4 did not (Figure 2C). As IFNγ and LPS are highly related to type 1 inflammation (i.e., M1 macrophages), this finding is in line with the idea that Sema3G - Nrp2 axis plays a pathogenic role in RA. We also sought to know if macrophages in human RA synovium express Nrp2. To address this issue, we re-analyzed the publicly available single-cell RNA-seq data of RA synovium [22]. Consistent with the data obtained from the CIA model, CD14-positive monocyte/macrophage population expressed Nrp2 (Figure 2D), but other immune cells such as T cells and B cells did not. Together, these results suggest that macrophages could respond to Sema3G in the inflamed joint.

**Figure 2.**
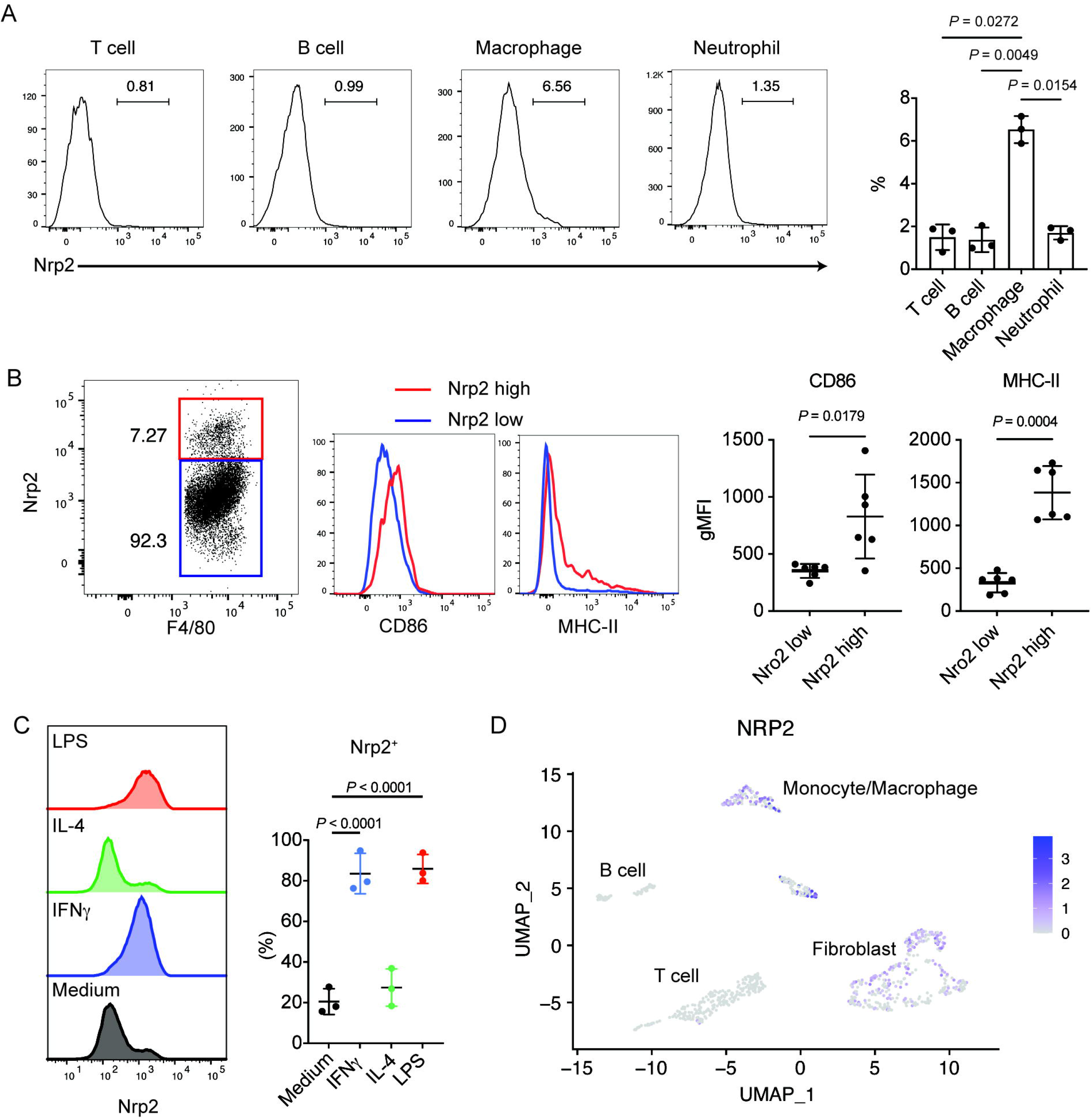
Nrp2 expression in activated macrophages during joint inflammation. **A.** Nrp2 expression on immune cells in CIA synovium. The hind paws of CIA-induced mice were digested and subjected to flow cytometric analysis. The representative histograms of Nrp2 staining and the cumulative data are shown. Data are expressed means ± SD. N = 3 from 3 independent experiments. **B.** The characteristics of Nrp2-positive macrophages. The representative histograms of CD86 and MHC class II expression on joint-infiltrating macrophages (left) and their cumulative data (right) are shown. N = 6 from 2 independent experiments. **C.** The expression of Nrp2 on BMMs. BMMs were cultured under indicated conditions, and Nrp2 expression was determined by flow cytometry. The representative histograms and the cumulative data are shown. N = 3 from 3 independent experiments. **D.** NRP2 expression in synovium-infiltrating cells in RA. The deposited single-cell RNA-seq data was re-analyzed, and several cell types were defined by UMAP. Expression of NRP2 is indicated with purple dots. The statistical analyses were performed using an unpaired t-test.

### Inflammatory arthritis models are mild in Sema3G^-/-^ mice

To address the roles of Sema3G during arthritis, Sema3G^-/-^ mice and their littermate control mice were subjected to CIA. As we found that Sema3G-heterozygous mice (Sema3G^+/-^ mice) expressed a comparable level of *Sema3g* in the spleen, kidney, and pancreas to wild-type mice (data not shown), we used Sema3G^+/-^ mice as controls. Notably, the clinical score was significantly lower in Sema3G^-/-^ mice than in control Sema3G^+/-^ mice (Figure 3A). Moreover, the pathological scores were lower in Sema3G^-/-^ mice (Figure 3B). We also measured IL-6 and TNFα in the sera to assess the global inflammatory conditions in these mice. While there was no difference in TNFα levels, serum IL-6 was significantly lower in Sema3G^-/-^ mice (Figure 3C).

**Figure 3.**
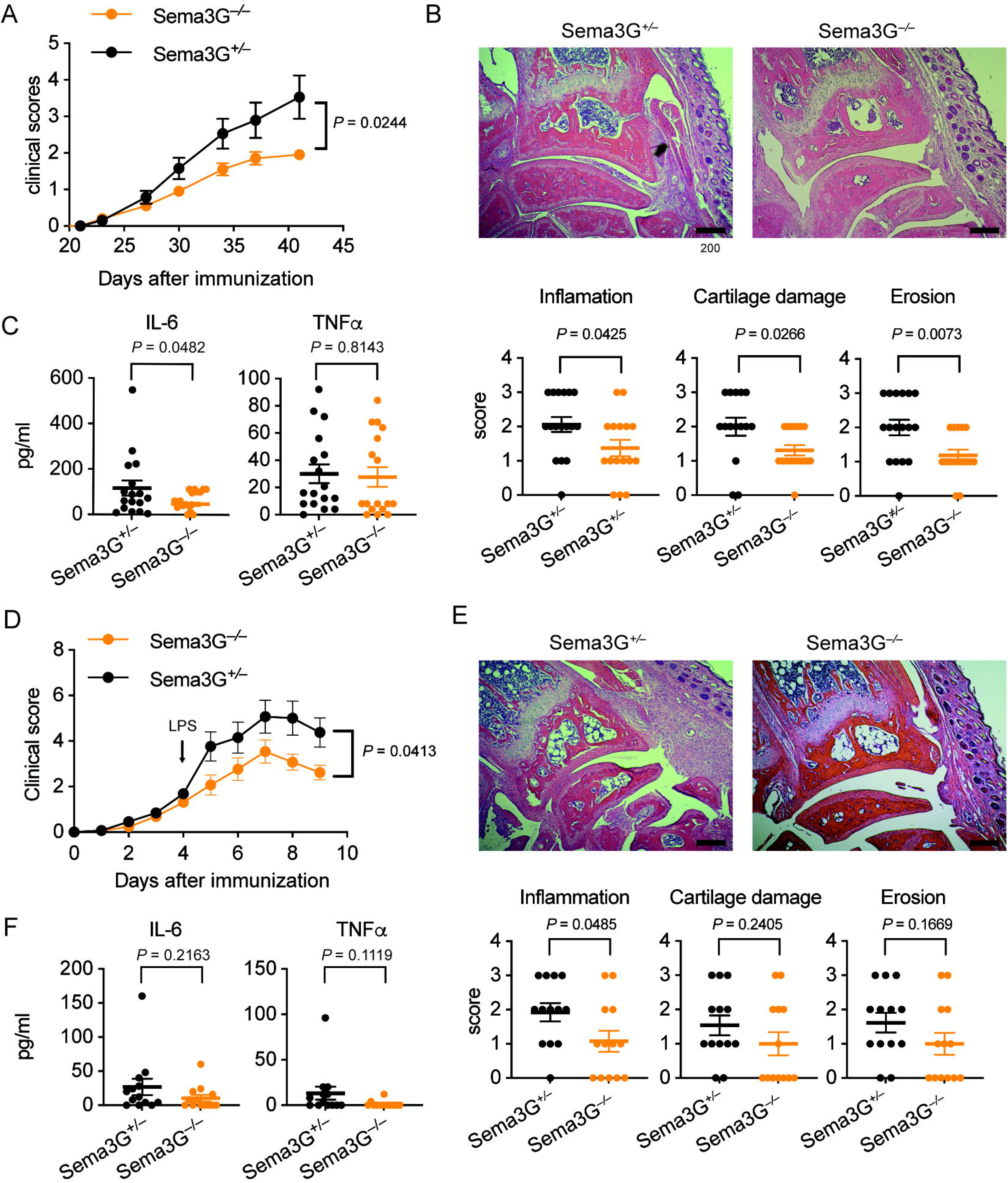
Attenuated joint inflammation in Sema3G-deficient mice. **A.** The clinical score of Sema3G-deficient (Sema3G^-/-^) mice and their littermate controls (Sema3G^+/-^) in CIA. Data were obtained from 4 independent experiments (N = 17 in each group). **B.** The pathological scores of CIA. The front paws were subjected to H&E staining, and the inflammation, the cartilage damage, and the erosion scores were separately assessed. **C.** The inflammatory cytokines in sera. The sera were collected on day 42 and subjected to ELISA to measure IL-6 and TNFα. **D.** The clinical score of Sema3G^-/-^ mice and Sema3G^+/-^ in CAIA. Data were obtained from 4 independent experiments (N = 13 in each group). **E.** The pathological scores of CAIA. The front paws were subjected to H&E staining, and the inflammation, the cartilage damage, and the erosion scores were separately assessed. **F.** The inflammatory cytokines in sera. The sera were collected on day 9 and subjected to ELISA to measure IL-6 and TNFα. Data are expressed as means ± SEM. For the analyses of the clinical score, 2-way ANOVA was used. For the pathological scores, an unpaired t-test was used.

It is well known that both innate and adaptive immune responses are required to develop CIA [23]. Since we next sought to dissect the mechanisms by which Sema3G deteriorates CIA, we employed CAIA in which adaptive immunity is less critical in the development of arthritis [24]. In this model, an anti-collagen antibody was injected intravenously into mice on day0, and LPS was injected intraperitoneally to boost inflammation on day4. We found that Sema3G^-/-^ mice showed attenuated clinical scores in CAIA when comparing overall disease courses with control Sema3G^+/-^ mice (Figure 3D). Of note, the deterioration rate was not remarkably changed after LPS stimulation in Sema3G^-/-^ mice, while control mice showed obvious responses to LPS stimulation (Figure 3D). Indeed, there was no statistical difference when comparing the clinical scores until day4 (*P* = 0.233). These findings suggest that Sema3G^-/-^ mice are hyporesponsive to LPS but normally respond to the inflammation-inducing antibody.

We next analyzed the histological score in CAIA. While there were no statistical differences in the cartridge damage and erosion score, the inflammatory score was significantly lower in Sema3G^-/-^ mice (Figure 3E). We also observed lower IL-6 and TNFα in the sera in Sema3G^-/-^ mice with no statistical significance (Figure 3F). Together, these results indicate that Sema3G deteriorates inflammatory arthritis and acts mainly on innate immunity.

### Sema3G promotes joint inflammation through macrophage migration and proliferation

We sought to know the mechanisms by which Sema3G deteriorates inflammatory arthritis. First, we performed a transwell migration assay on LPS-stimulated BMMs to analyze the chemotactic property of Sema3G because semaphorins are best characterized as neuron guidance factors. Interestingly, we found that Sema3G promoted macrophage migration to some extent, but the chemotactic property was milder than MCP1 (Figure 4A).

**Figure 4.**
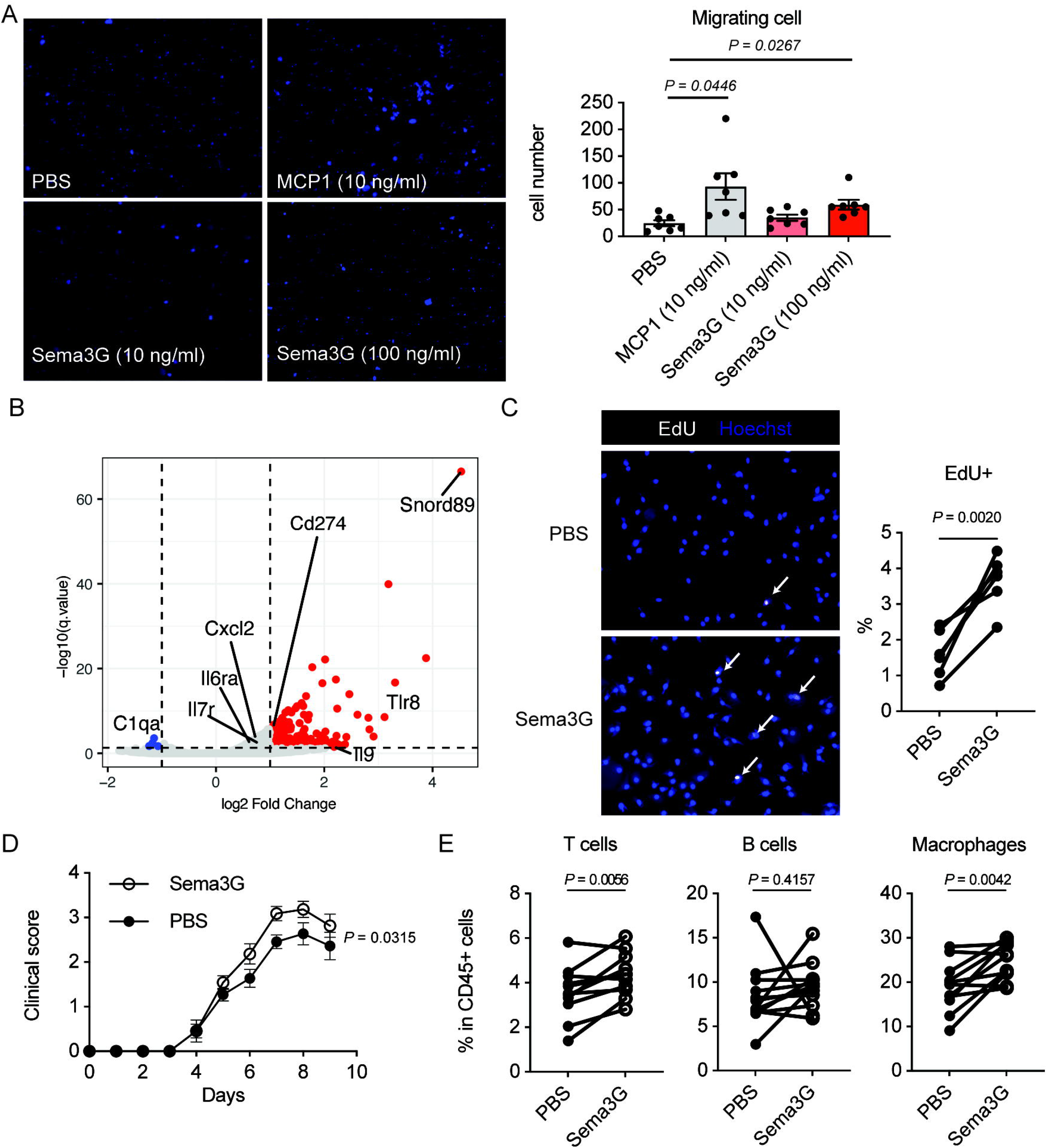
Enhanced macrophage proliferation by Sema3G. **A.** Chemotaxis assay. LPS-stimulated BMMs were subjected to a transwell migration assay. PBS (negative control), MCP1 (positive control), or various concentrations of Sema3G were added to the lower chambers. Cell migration towered to the lower chamber was determined by Hoechst staining. The pictures were taken for 5 different areas, and Hoechst-positive round cell numbers were counted. N = 7 from 3 independent experiments. One-way ANOVA test followed by Dunnet test was used for the statical analysis. **B.** The volcano plot of Sema3G-stimulated BMMs. Genes differentially expressed are plotted as red dots (upregulated by Sema3G) or blue dots (downregulated by Sema3G). Some immune-related gene names are labeled. **C**. EdU uptake in Sema3G-stimulated BMMs. Sema3G-stimulated or control (PBS) BMMs were pulsed with 10 μM EdU for 2 hours, and EdU and Hoechst were detected by immunofluorescent analysis. The pictures were taken for 10 different areas, and the percentage of EdU-positive cells over Hoechst-positive cells was calculated. Data were obtained from 3 independent experiments. N = 6. **D.** The clinical score of Sema3G- or PBS-injected paw during CAIA. WT mice were subjected to CAIA. Sema3G (100 ng) was injected into the right footpad, and PBS was injected into the left footpad daily from day3 to day9. N = 11 from 2 independent experiments. **E.** Infiltrating cell subsets to each paw on day 9 of CAIA. The hind paws were digested, and cells recovered were analyzed by flow cytometry. The percentages of T cells, B cells, and macrophages are shown. A paired t-test was used for the statistical analyses.

Next, we performed the transcriptome analysis of Sema3G-stimulated BMMs to understand further the molecular mechanisms by which Sema3G drives joint inflammation. In this experiment, LPS-stimulated BMMs were further cultured with or without recombinant Sema3G for 18 hours and subjected to RNA-seq analysis. Many genes were upregulated by Sema3G stimulation, and among these genes, *Snord89* was the transcript highly expressed in Sema3G-stimulated BMMs (Figure 4B). Snord89 is a non-coding RNA, and it has been reported that Snord89 is related to cell proliferation through Myc activation [25]. We, therefore, hypothesized that Sema3G upregulates Snord89 to proliferate macrophages, resulting in deteriorating arthritis. To clarify this idea, we analyzed macrophage proliferation upon Sema3G stimulation. In this experiment, LPS-stimulated BMMs were cultured in the presence or absence of Sema3G for 48 hours. Since BMMs adhered to plastic surfaces very firmly and it was difficult to count cell numbers with a hemocytometer, BMMs were incubated with EdU for the last 2 hours, and de novo DNA synthesis was evaluated by measuring the incorporation of EdU. As shown in Figure 4B, the percentage of EdU-positive cells among Hoechst-positive cells was increased in Sema3G-stimulated BMMs compared to that in PBS-stimulated BMMs (Figure 4C), indicating that Sema3G promotes cell proliferation. Collectively, enhanced macrophage proliferation by Sema3G may be a mechanism to accelerate joint inflammation.

Finally, we assessed whether the local administration of Sema3G affects the severity of arthritis and macrophage migration/proliferation in vivo. CAIA was induced in wild-type mice, and 100 ng recombinant Sema3G (10 μl) was injected into the footpad of the left side daily. As a control, 10μl PBS was injected into the footpad of the right side daily. Consistent with the in vitro data, Sema3G-injected footpads showed a higher clinical score than PBS-injected ones (Figure 4D). In addition, the percentage of macrophages was significantly higher in Sema3G-injected left footpad than in PBS-injected right footpad (Figure 4E). The frequency of T cells was also increased in the Sema3G-injected footpad (Figure 4E). These results suggest that Sema3G promotes macrophages migration/proliferation in vivo and in vitro.

## Discussion

In this study, we show the role of Sema3G in the pathogenesis of inflammatory arthritis. We identified Sema3G as one of the downregulated genes by MTX treatment in RA patients. We also found that Sema3G was expressed in the inflamed joint in humans and mice. Activated macrophages expressed Nrp2, the receptor for Sema3G. In vivo, Sema3G^-/-^ mice displayed mild disease in both CIA and CAIA, and Sema3G administration deteriorated arthritis in CAIA. Mechanistically, Sema3G promoted the migration and proliferation of macrophages.

Semaphorins were first identified as neural guidance proteins [26]. Semaphorins are largely classified into 8 classes; only class 3 semaphorins are secreted, whereas the rest of the classes are transmembrane proteins in mammals [27]. Recent studies have revealed their crucial roles in cardiovascular growth [28], bone homeostasis [29], and immune responses [20, 30]. Sema3G was initially defined as a repulsive factor of sympathetic axons [21] and is now related to several physiological and pathological processes [18, 31–33]. We previously reported the protective role of Sema3G in LPS-induced kidney injury [33]. In podocytes, Sema3G suppresses the production of inflammatory cytokines/chemokines such as IL-6 and CCL2 upon LPS stimulation. In this study, we performed unbiased RNA-seq analysis of Sema3G-stimulated BMMs and found no difference in *IL6* level (data not shown). Thus, it is plausible that Sema3G has cell-type-specific functions during inflammation. Further studies are required to understand how Sema3G affects immune responses in each cell type.

Macrophages have various roles in the pathogenesis of inflammatory arthritis. Macrophages are classified into two subsets, namely M1 and M2 macrophages. M1 macrophages are thought to be pro-inflammatory and implicated in the pathogenesis of autoimmune diseases, whereas M2 macrophages have anti-inflammatory potential [34]. In human RA, macrophages are one of the most abundant immune cell types that infiltrate the synovium. It has been reported that RA synovial fluid contains more M1 macrophages than M2 macrophages [35]. In addition, several studies have revealed that the number of M1 macrophages correlates with disease activity and joint damage [35, 36]. Like what is observed in humans, M1 macrophages have pathogenic roles in the murine arthritis models [37–39]. These findings suggest that M1 macrophages are highly implicated in the pathogenesis of RA.

In this regard, we showed that Nrp2 expression on BMMs was significantly elevated upon LPS or IFNγ stimulation (Figure. 2C), which favors M1 macrophage differentiation. Furthermore, Nrp2-positive macrophages expressed M1 markers in the synovium (Figure 2B and 2C). These activation markers are typically expressed in M1 macrophages so that Nrp2-positive macrophages may have an M1 phenotype. On the other hand, M2 macrophage-skewing cytokine IL-4 did not promote Nrp2 expression. These findings indicate that M1-like pro-inflammatory macrophages preferentially respond to Sema3G and proliferate in the inflamed synovium, suggesting that Sema3G is the pro-inflammatory secreting protein in the arthritic joint.

To our interest, Sema3G^-/-^ mice were resistant to LPS stimulation, although Sema3G^-/-^ mice showed normal initial response to anti-collagen antibody during CAIA (Figure 3D). It has been reported that the activation of alternative pathway of complement accompanied with IgG-Fc receptor coupling is vital for the induction of arthritis [40]. Anti-collagen antibodies are deposited on the cartridge surface and recognized by FcRγIII-bearing cells such as neutrophils, mast cells, and macrophages. Activation of the antibody-FcRγIII and complement-C5aR pathways promotes the production of inflammatory cytokines such as IL-6 and TNFα, resulting in joint inflammation. Interestingly, RNA-seq analysis revealed that Sema3G stimulation increased the expression of neither *FcRγIII* nor *C5aR* in BMMs (data not shown). So, we speculate that Sema3G has a minimal role in antibody-mediated immune responses but directly affects macrophage migration and proliferation. This feature may be beneficial when considering Sema3G as a therapeutic target because antibody-mediated immune responses are essential for pathogen clearance [41].

This study has several limitations. First, although we identified Sema3G as a molecule whose expression was down-regulated by MTX treatment, we could not address the molecular mechanism underlying MTX-mediated suppression of Sema3G expression because of the difficulty of in vitro experiments using MTX. Second, Sema3G-mediated immune regulation during arthritis is not fully addressed. While we found Nrp2 was expressed on activated macrophages, some lymphoid cells and neutrophils were also positive for Nrp2 in the synovium. Several studies have revealed the crucial roles of semaphorin family members in T cell differentiation and activation [42–47]. Indeed, we found that T cells were increased in the inflamed joint upon local Sema3G administration (Figure 4D). Thus, it is possible that the Sema3G-Nrp2 axis also has roles in T cell-mediated immune responses during arthritis. Third, we could not assess the therapeutic potential of Sema3G neutralization directly. Although the severity of CIA and CAIA was reduced in Sema3G^-/-^ mice, antibody-mediated neutralization of Sema3G after the initiation of arthritis would be more important to know whether Sema3G could be a therapeutic target. Further studies are required to address this point when a neutralization antibody against Sema3G is available.

In conclusion, Sema3G is implicated in the pathogenesis of RA, and the neutralization of Sema3G may lead to the development of a novel therapy belonging to a new category.

## Acknowledgment

We thank Ms. Kazumi Nemoto and Maho Yoshino for the technical support and all the members of Nakajima Lab for valuable discussion. Dr. Shigeo Hagiwara helped to recruit patients who received joint surgery.

## Author contribution

**Study conception and design.** Shoda, Tanaka, Nakajima.

**Acquisition of data.** Shoda, Tanaka, Etori, Hattori, Kasuya, Ikeda, Yuko Maezawa, Suto, Suzuki, Nakamura, Yoshiro Maezawa.

**Analysis and interpretation of data.** Shoda, Tanaka, Takemoto, Betsholtz, Yokote, Ohtori, Nakajima.

